# Oligodendrocytes in the mouse optic nerve originate in the preoptic area

**DOI:** 10.1101/078774

**Authors:** Katsuhiko Ono, Kengo Yoshii, Hiroyuki Tominaga, Hitoshi Gotoh, Tadashi Nomura, Hirohide Takebayashi, Kazuhiro Ikenaka

## Abstract

The present study aims to examine the origin of oligodendrocyte progenitor cells (OPCs) in the mouse optic nerve (ON) by labeling OPCs in the fetal forebrain. The labeling of OPCs in the ON was performed by injection of a retrovirus vector carrying the *lacZ* gene into the lateral ventricle, or by inducible Cre/loxP of Olig2-positive cells. The retrovirus labeling revealed that ventricular zone-derived cells of the fetal forebrain relocated to the ON and differentiated into oligodendrocytes. In addition, lineage tracing of Olig2-positive cells and whole mount staining of PDGFRα-positive cells demonstrated that OPCs appeared by E12.5 in the preoptic area, and spread caudally to enter the ON. Our results also suggest that OPCs generated during the early stage are depleted from the ON after maturation.

## Introduction

Mechanisms underlying oligodendrocyte (OL) development have been uncovered by extensive studies after the isolation of O-2A (oligodendrocyte-type 2 astrocyte) progenitor cells from newborn rat optic nerves (ON) (Raff et al., 1984). One of the most remarkable findings is that PDGF is a strong mitogen for O-2A cells and that O-2A cells express PDGF receptor alpha (PDGFRα) (Richardson et al., 1988). O-2A cells differentiate into OLs when grafted into the forebrain of newborns (Espinosa de los Monteros et al., 1993), indicating that O-2A progenitors correspond to oligodendrocyte progenitor cells (OPCs) *in vivo*. PDGFRα is the first reliable lineage marker for OPC *in vivo* (Pringle et al., 1992). During the past quarter century, OPC development has been extensively studied in the vertebrate central nervous system (Miller, 1996; Miller, 2005; Naruse et al., 2016; Ono et al., 1995; Ono et al., 1997; Orentas and Miller, 1998; Richardson et al., 2000). Although OPCs in the ON are considered to originate in the basal forebrain (Garcion et al., 2001; Small et al., 1987), the precise origin of OPCs in the rodent ON has not been examined experimentally.

The present study aims to elucidate the origin of OPCs in the ON in the developing mouse. We used 2 strategies to label OPCs in the basal forebrain; the first is labeling mitotic cells in the fetal ventricular surface by an injection of high-titer retrovirus vector carrying the *lacZ* gene into the fetal lateral ventricle (Nanmoku et al., 2003), and the second is to label Olig2-positive progenitor cells with EGFP by ligand-inducible Cre/loxP (Masahira et al., 2006). The results suggest that OPCs in the ON arise in the preoptic area (POA) by E12.5, relocate caudally to enter the ON, and differentiate to myelinating OLs, but not to GFAP-positive astrocytes. In addition, our results also suggest that OPCs generated in the early stage are depleted from the ON after maturation.

## Results and Discussion

### Cells derived from forebrain ventricular zone migrate to the ON

To examine whether forebrain ventricular zone (VZ)-derived cells at fetal stages migrate into the ON, we labeled VZ cells with high-titer retrovirus vector carrying the *lacZ* gene. When the retrovirus was injected into the fetal lateral ventricle at E12.5, E14.5, or E15.5, a considerable number of the VZ cells were labeled with LacZ 2 days after the injection. In later stages, a great number of LacZ+ cells were observed in the forebrain (Nanmoku et al., 2003). Among these cases, 2 ONs after the E14.5 injection and 2 after the E15.5 injection contained a great number of LacZ-positive cells; nearly 2,000 labeled cells were observed at P30 (n=2 ONs) and approximately 800 cells at P180 (n=2 ONs) (Supplementary Table 1). No labeling was observed in the ON following the E12.5 injection. In the P30 ON, the labeled cells were densely distributed (Fig. 1A), and, therefore, individual cell morphology was not clear (Fig. 1B). In the P180 ON, nearly all, if not all, of the LacZ-positive cells had a cell body with several processes extended in parallel to the longitudinal axis of the ON (Fig. 1C), which is a typical myelinating OL shape. Astrocyte-like cells with a star shape were rarely detected in the ON (not shown). This result demonstrate that forebrain VZ-derived cells at these fetal stages relocate to the ON and differentiate into OLs.

**Figure 1.**
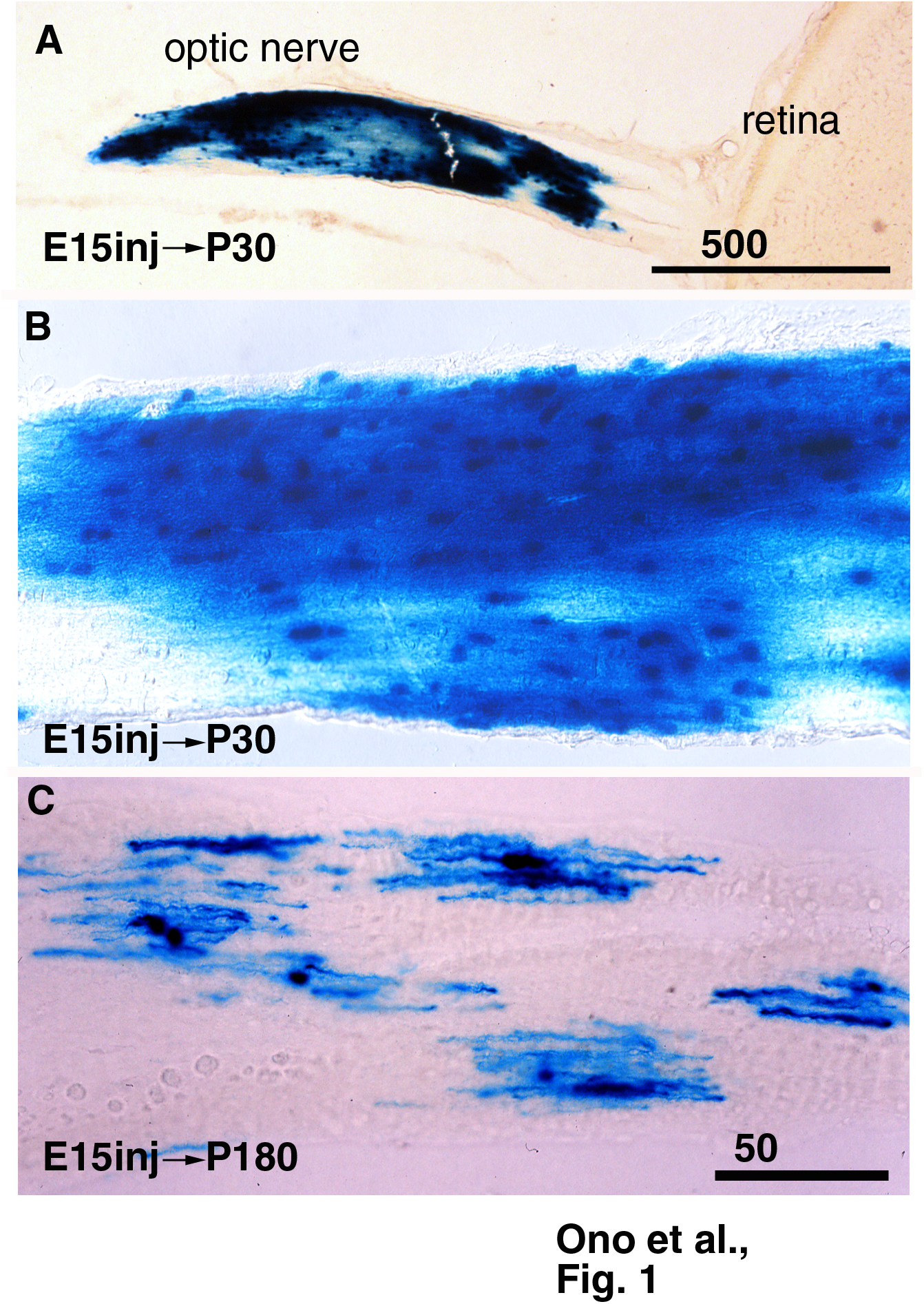
LacZ-positive cells in the adult ON following retroviral injection into the fetal lateral ventricle. (A) A low magnification picture of the P30 ON following E15.5 viral injection. (B) A high magnification picture of the P30 ON. (C) LacZ-positive cells in the P180 ON following E15.5 viral injection. The labeled cells show profiles of oligodendrocytes.

### OPCs for ON may arise in the preoptic area

We next examined the distribution of OPCs in the fetal basal forebrain close to the ON with Olig2 and PDGFRα as markers. Olig2-positive cells were observed in the VZ of the POA and caudal hypothalamus, but not in the VZ in the level of the optic stalk at E12.5 (Supplementary Fig. 1). Whole mount staining of PDGFRα demonstrated that OPCs in the ventral diencephalon were initially localized in the anterior part of the POA at E13.5 (Fig. 2AC). Subsequently, PDGFRα-positive cells dispersed caudally to the rostral and caudal hypothalamus including the wall of the optic recess (Fig. 2BD). Coronal sections of E12.5-E14.5 basal forebrain and hypothalamus also showed a rostral-to-caudal gradient of distribution of PDGFRα-positive OPCs and the POA contained the labeled cells most abundantly (Supplementary Fig. 2). The optic stalk is devoid of OPCs at these stages (supplementary Figs. 1GK and 2ILM).

**Figure 2.**
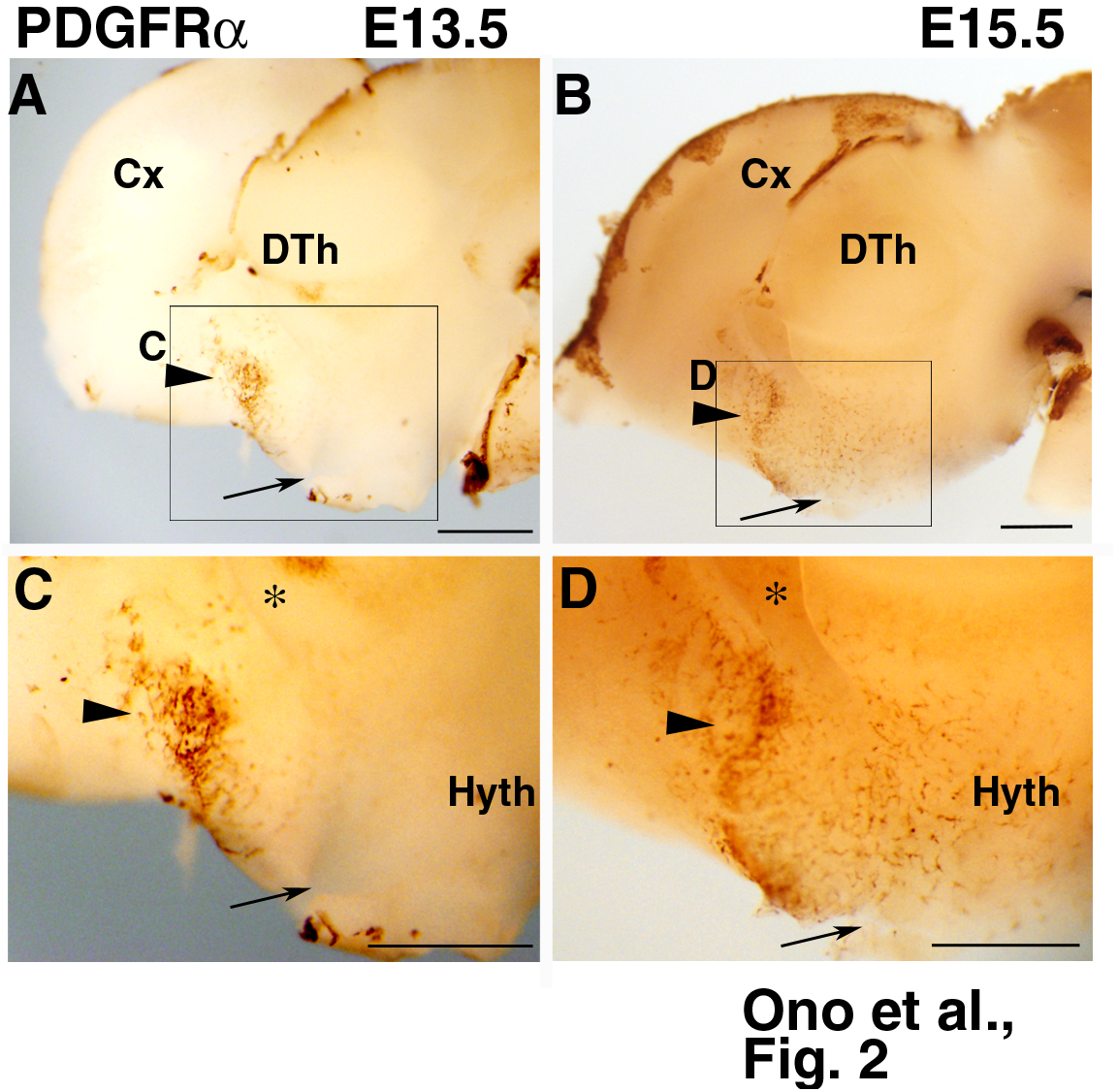
Initial appearance and dispersal of OPCs in the hypothalamus. Whole-mount immunohistochemistry of PDGFRα. Medial view of the brains cut along the midline. Boxed areas in A and B are magnified in C and D, respectively. Arrows indicate the optic recess. (A) E13.5. PDGFRα-positive cells are observed in the rostral part of the preoptic area (arrowhead). (B) E15.5. PDGFRα-positive cells are widely distributed in the preoptic area (arrowhead), and scattered caudally beyond the optic recess (arrow). Asterisks in C and D indicate interventricular foramen. Cx, cerebral cortex. DTh, dorsal thalamus, Hyth, hypothalamus. Bars= 500 µm.

We then performed short-term lineage tracing of Olig2-positive cells to examine when OPCs appeared in the ON. TM was injected into double-heterozygous animals at E11.5 or at E11.5+E12.5, and E17.5 ONs were observed after double-immunostaining for EGFP and PDGFRα. The results demonstrated that EGFP/PDGFRα-double positive cells appeared in the ON treated with TM at E11.5+E12.5 (n=4 ONs, 2 mice; Supplementary Fig. 3AB; more than 70% of EGFP-positive cells expressed PDGFRα), while the ON at E11.5 did not contain EGFP-positive cells (n=6, 3 mice; not shown), indicating that OPCs appeared in the ON in the forebrain by E12.5.

We then performed short-term lineage tracing with a 1- or 2-day survival period after TM treatment to observe the origin of OPCs retrospectively. In the E13.5 basal forebrain after E12.5 TM treatment, EGFP-positive cells in the POA expressed PDGFRα (Fig. 3A-D), while EGFP-positive cells in the caudal hypothalamus were not positive for PDGFRα (Fig. 3EFG). In the E15.5 or older forebrain, 3 or more days after TM treatment, EGFP/PDGFRα double-positive cells were observed to enter the optic chiasm from the dorsally adjacent neuroepithelium (n=6 mice, Fig. 3GH).

**Figure 3.**
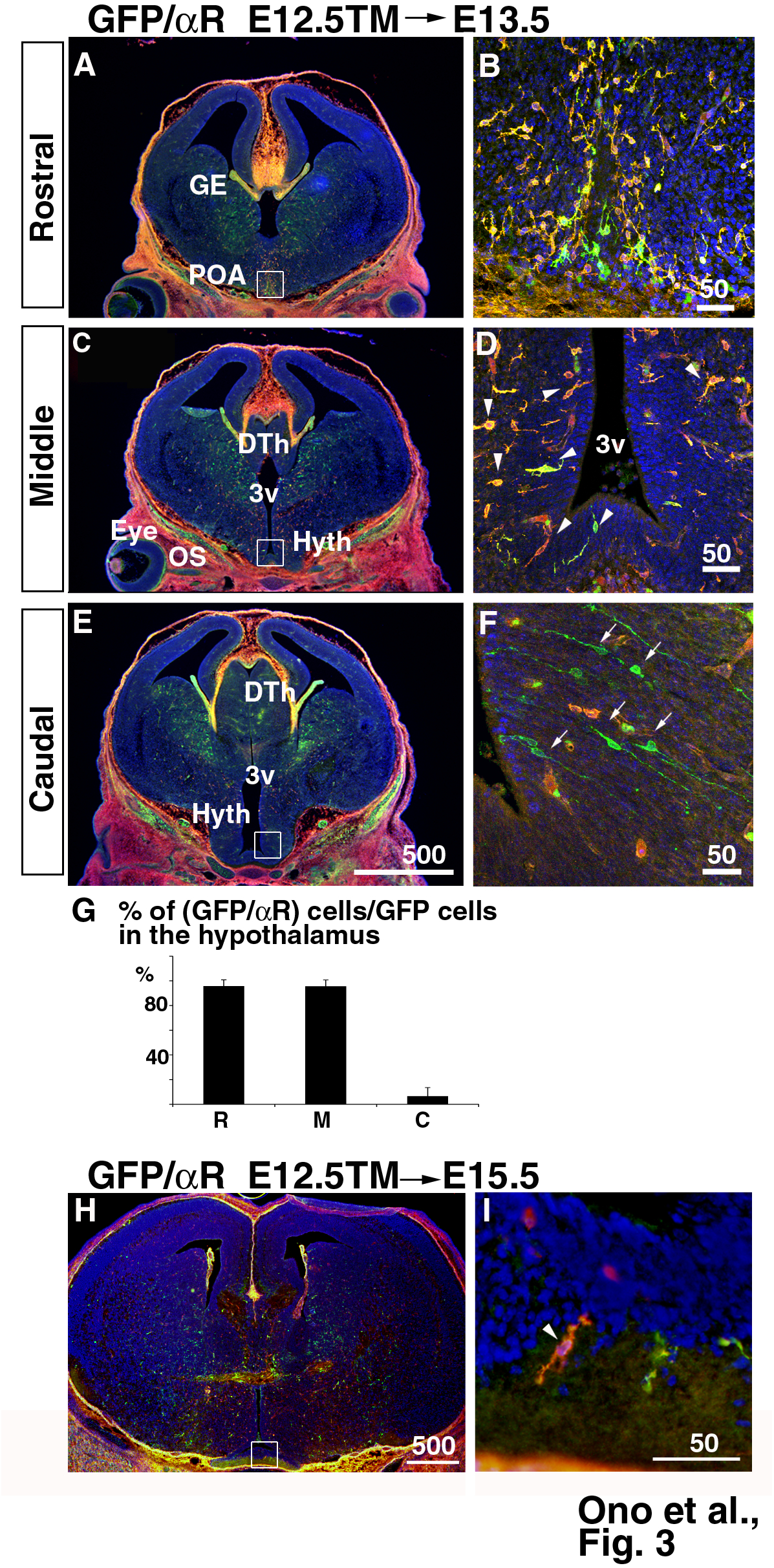
Olig2 lineage PDGFRα-positive cells in the basal forebrain and the ON at fetal stages. (A-F) EGFP-positive cells in the E13.4 basal forebrain following E12.5 TM injection. Nearly all EGFP-positive cells in the preoptic area (POA) and rostral hypothalamus (Hyth) are PDGFRα-positive (A-D; arrowheads), while EGFP-positive cells in the caudal hypothalamus are not (arrows). (G) The percentage of EGFP/PDGFRα double-positive cells in the EGFP-positive cell population in the hypothalamus. Sections were grouped as the anterior (A), middle (C), and most-caudal (E) levels (n=3 mice). (H, I) EGFP/PDGFRα double-positive cells (arrowhead) are observed to enter the optic chiasm from the overlying neuroepithelium at E15.5. Boxed areas in the left columns are magnified in the right columns. Bars in B, D, F, and I = 50 µm; in E, H = 500 µm

### Olig2-positive cells in the E12.5 forebrain develop into myelinating OLs in the ON

We next performed long-term lineage tracing of fetal Olig2-positive OPCs in the forebrain to see whether they differentiated into myelinating OLs in the ON. Double-heterozygous animals treated with TM at E12.5 were born and grew to P30 (n=6 mice) or P180 (n=3 mice). In both stages (Fig. 4), EGFP-positive cells and processes formed clusters, which were localized intermittently in the ON in both stages (Fig. 4A-C). The majority of EGFP-positive cells were CC1-positive, and thus mature OLs (Fig. 4ACG). EGFP-positive processes were extended in parallel to axon arrangement in longitudinal sections, and were positive for proteolipid protein (PLP), a myelin protein (Fig. 4J). A small fraction of EGFP-positive cells were PDGFRα-positive in the P30 ON while no or few double-positive cells were detected at P180 (Fig. 4DEH). We estimated the total number of EGFP-positive cells in the ON. Approximately 800 cells were found in the P30 ON (n=6), although there was a large individual variation from 80 to 2,000, while nearly 400 cells per ON were detected at P180 (Fig. 4F). In the P30 ON, 80% of EGFP-positive cells were CC1-positive mature OLs (Fig. 4A) and 11% of them were PDGFRα-positive OPCs (Fig. 4BGH). In contrast, almost all EGFP-positive cells were CC1-positive mature OLs in the P180 ON, as above-mentioned (Fig. 4GH).

**Figure 4.**
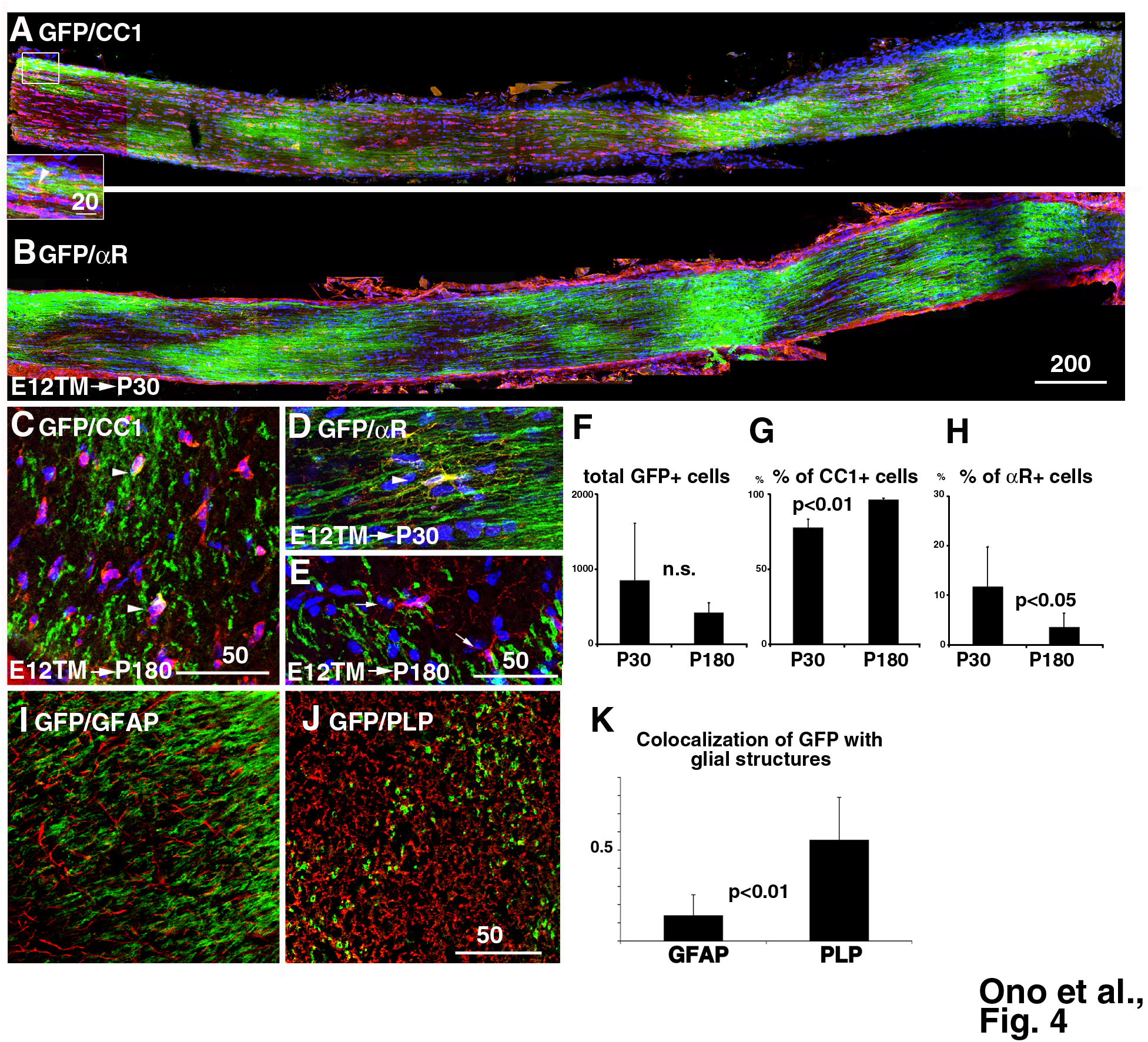
Appearance of Olig2-lineage cells in the adult ON following E12.5 TM injection. (A, B) Longitudinal sections of the P30 ON, double-stained with anti-EGFP and anti-CC1 (A; OLs) or anti-PDGFRα (B; OPCs) antibodies. Boxed area in C is magnified in the inset. (E) EGFP/CC1 double-positive cells in the coronal section of the P180 ON. Arrowheads indicate double-positive cells. (F) Appearance of EGFP/PDGFRα double-positive OPCs in the P30 ON. (G) No or few EGFP/PDGFRα double-positive OPCs in the P180 ON. (H) Estimation of the total EGFP-positive cells in single ONs at P30 (n=6 ONs) and P180 (n=6). (I) Percentage of EGFP/CC1 double-positive cells among the EGFP-positive cell population. (n=6 in P30 and P180). (J) Percentage of EGFP/PDGFRα double-positive cells among the EGFP-positive cell population. (n=4 in P30 and n=6 in P180). (K-M) Colocaliztion analysis of EGFP-positive structures with glial structures in the P30 ON. EGFP-positive structures colocalized with PLP (L and M) significantly more than with GFAP (K). Bar in D = 200 µm; in A, C, E, J = 50 µm; in B = 25 µm.

We also stained the ON with anti-EGFP and anti-GFAP antibodies to examine whether fetal Olig2-positive cells develop into astrocytes. No or few overlapping was noticed in the double-stained sections from both P30 and P180 animals. Colocalization analysis in single optical sections from the P30 ON demonstrated that EGFP-positive structures significantly overlapped more often with PLP than with GFAP-positive structures (Fig. 4I-K). These results indicate that fetal forebrain Olig2-positive cells relocate in the ON and develop into mature myelinating OLs, but not astrocytes, and that early generated OPCs were depleted by the adult stage.

The present retrovirus injection study clearly demonstrated that OLs in the ON are derived from forebrain VZ cells in the fetal stage. In addition, whole-mount immunostaining of PDGFRα revealed that OPCs in the hypothalamic area were initially restricted to the POA, and therefore, the POA is the site of OPC origin in the ON. We previously demonstrated in chick embryos that OPCs arise from the floor of the third ventricle adjacent to the optic chiasm (Ono et al., 1997). In the present study, EGFP/PDGFRα double-positive cells were observed in the chiasmal region at E15.5. However, no Olig2-positive cells were observed in the future chiasmal region at E12.5 (Supplementary Fig. 1) when TM treatment was performed. Short-term lineage tracing demonstrated that double-positive cells were most abundant in the POA, whereas none of the EGFP-positive cells were PDGFRα-positive in the caudal hypothalamus. Olig2-positive cells in the caudal hypothalamus co-express neurogenin-2 (Ono et al., 2009), and therefore these cells probably differentiate into neurons, but not into OPCs. These results strongly indicate that chiasmal double-positive cells are derived from the POA. Together, these results suggest that OPCs in the ON arise in the anterior part of the POA (Supplementary Fig. 4). In the late fetal stages, Olig2-positive cells are distributed not only in the POA, but also the rostral and caudal hypothalamus (Ono et al., 2008). Such Olig2 cells could also generate OPCs in the hypothalamus and the ON, and will need to be uncovered by future analysis.

O-2A progenitor cells *in vitro* differentiate into either type 2 astrocytes or into OLs, whereas they differentiate into OLs when they are grafted to the forebrain of newborns (Espinosa de los Monteros et al., 1993; Raff et al., 1984). In the present study, most, if not all, VZ-derived cells labeled with LacZ differentiated to OLs in the ON (Fig. 1C), while astrocyte-like cells were rarely detected in the ON. In addition, Olig2-positive cells which migrated to the ON from the forebrain were shown to develop only into OLs. Although colocalization analysis showed that 10% of EGFP-positive structures overlapped with GFAP-positive structures, we did not observe overlapping distribution of EGFP with GFAP in the ON. Only PLP- or CC1-positive structures were clearly colocalized with EGFP-positive structures. Therefore, the present result indicates that OPCs *in vivo* do not differentiate into astrocytes in the ON. Olig2-positive progenitor cells have been reported to differentiate into astrocytes as well as OLs in other regions of the CNS (Ono et al., 2008; Tatsumi et al., 2008), indicating that there might be inhibitory factors in the ON for OPCs to develop to astrocytes, which are specifically localized in the ON.

In conclusion, the present study demonstrated that OPC in the ON arise in the POA of the developing mouse brain by E12.5, probably under the influence of local signals, such as sonic hedgehog (Orentas et al., 1999), expressed in the POA (Ono et al., 2008). They gradually disperse caudally to enter the ON from the chiasmal region. There may be species difference between the mouse and the chick with respect to the origin of early OPCs in the ON (mouse, present study; chick, Ono et al., 1997). A phylogenic comparative study would elucidate the significance of species differences.

## Materials and methods

### Animals and tissue preparation

The animals used in this study were Olig2^KICreER^ (Takebayashi et al., 2002), ROSA26-GAP43-EGFP (Nakahira et al., 2006), and wild-type ICR mice (Slc, Shizuoka, Japan). The day when the vaginal plug was detected was regarded as E0.5. Genotyping was performed as previously described (Takebayashi et al., 2002; Tatsumi et al., 2008). All animal experiment procedures were approved by the Animal Research Committee of Kyoto Prefectural University of Medicine and of National Institute for Physiological Sciences.

To label Olig2 lineage cells, Olig2^KICreER^;ROSA26-GAP43-EGFP double heterozygous mice were mated with wild-type mice, and tamoxifen (TM; 3mg/animal) was intraperitoneally injected into pregnant mothers with fetuses at E11.5 or E11.5 and E12.5 (E11.5+E12.5), E12.5, E14.5, or E15.5, as previously described (Masahira et al., 2006). Pregnant mice were deeply anesthetized with pentobarbital (100 mg/kg body weight), and fetal mice were removed from the uterus. Fetal mice brains at E15.5 or younger were fixed by immersion in 4% paraformaldehyde (PFA) in phosphate-buffered saline (PBS) overnight. Mouse brains at E17.5 or after birth were fixed by perfusion with 4% PFA through the heart, and the ON was isolated and immersed in 20% sucrose in PBS. The ON was cut transversely or longitudinally with a cryostat at 20μm in thickness, and sections were thaw-mounted onto MAS-coated glass slides (Matsunami Glass Co., Tokyo, Japan). Nomenclature of fetal brain structures is according to that used by Bulfone et al. (Bulfone et al., 1993) and by Nieuwenhuys et al. (Nieuwenhuys et al., 2008).

### Production and injection of high-titer retrovirus vector

Preparation of a retrovirus vector carrying the *lacZ* gene and injection of the vector into the fetal lateral ventricle were performed as previously reported (Nanmoku et al., 2003). We used the ON of the same samples as the previous report. The ON was cut with a vibratome at 100μm in thickness and stained with X-Gal as previously reported (Nanmoku et al., 2003).

### Immunohistochemistry and *in situ* hybridization

Immunohistochemistry was carried out as previously described (Ono et al., 2014). Primary antibodies used are listed in Supplementary table 1. *In situ* hybridization of *PDGFR*α was performed as previously described (Ivanova et al., 2003). Stained sections were observed under an epifluorescent or bright field microscope (BX-51; Olympus, Tokyo, Japan), or with a confocal laser scanning microscope (CLSM; FV-1000; Olympus).

Whole-mount immunohistochemistry of PDGFRα was performed in E13.5~E15.5 brains that were cut along the midline. They were treated with 0.1% H_2_O_2_ for 4 hours, with 2% BSA and 5% dimethylsulfoxyde for 5 hours, and subsequently incubated with primary antibody (1:1000) overnight. They were washed 3 times for 4 hours, and then incubated with biotinylated anti-goat antibody (1:1000) for 5 hours. After washing, they were processed with an ABC elite kit, color was developed by incubation with tris-HCl (50mM, pH7.4) containing diaminobenzidine and H_2_O_2_, and they were observed under a dissection microscope (SZX7, Olympus). Olig2-KO brains were used as a negative control.

### Image analysis

Colocalization analysis was performed with Fiji/ImageJ software (NIH). Dual color images of EGFP and GFAP, or of EGFP and PLP, were captured with a FV1000 CLSM. Manders split coefficients (Manders et al., 1993) above the threshold (tM) were calculated, which indicated, proportional to the amount of fluorescence, the colocalizing pixels in each color channel. Five regions from 1 image were averaged, and at least 3 images were analyzed and compared between GFAP and PLP.

## Acknowledgement

We would like to express our sincere appreciation to Ms. Masako Kawano for technical help, and to Dr. Martin Goulding for providing a reporter mouse. This work was supported by a Grant-in-Aid for Scientific research provided by MEXT, and also by a collaboration Grant provided by NIPS.

